# Sub-therapeutic Concentrations of Antibiotics Induce Prophage-driven Superinfection Exclusion and Fitness Cost in *Pseudomonas aeruginosa*

**DOI:** 10.1101/2024.11.20.624585

**Authors:** Michael J. Bucher, Cristian P. Puente, Naveen Sehdev, Daniel M. Czyż

**Affiliations:** Department of Microbiology and Cell Science, University of Florida, Gainesville, Florida, USA

**Keywords:** Bacteriophage, Pseudomonas aeruginosa, superinfection exclusion, antibiotics, fitness

## Abstract

Bacteria and other microbes can naturally produce antibiotics within their native soil environment, but often at sub-inhibitory concentrations; consequently, the exact role of antibiotics within bacterial native communities remains unknown. We have shown that subtherapeutic quantities of naturally occurring antibiotics can induce the *Pseudomonas* prophage Pf4, and superinfection of *Pseudomonas aeruginosa* cells by this phage leads to their reduced virulence, as demonstrated by impaired twitching motility, compromised macrophage evasion, and increased killing by macrophages *in vitro*. Thus, the production of subtherapeutic concentrations of antibiotics by environmental microbes may provide the producers an evolutionary advantage associated with reduced fitness induced by prophages in the competing bacteria. Collectively, these results reveal the role of naturally occurring antibiotics in altering fitness by phage-mediated superinfection exclusion and provide potential clinical implications in the application of phage therapy.

**Significance Statement:** This study provides insights into the mechanisms by which sub-inhibitory concentrations of environmentally-produced antibiotics induce *Pseudomonas aeruginosa* prophages, revealing a potential evolutionary strategy for competitive advantage among bacteria. By activating prophages, antibiotics can induce fitness defects in neighboring bacteria, impacting their motility, phagocytosis, and survival within macrophages. Such previously unrecognized role for environmental antibiotics in bacterial ecosystems may offer insights into enhancing phage therapy by exploiting phage-antibiotic synergies. Understanding phage-host interactions can enhance therapeutic strategies to mitigate bacterial infections and antimicrobial resistance.

## Introduction

Antibiotics, at sub-therapeutic concentrations, can trigger integrated prophages to transition from a lysogenic to a lytic state (1, 2). These antibiotics can be produced and released by many species of bacteria or other diverse microbes in the environment. Often, the concentrations of antibiotics released are too low to directly kill neighboring microbes due to the metabolic costs of producing these complex secondary metabolites, and their exact function at sub-inhibitory concentrations remains largely unknown (3–5). However, antibiotic-mediated activation of prophages may lead to bacterial killing through cell lysis, or impose fitness costs on the competing species, allowing the antibiotic producer to outcompete for resources (6).

*Pseudomonas aeruginosa*, a ubiquitous opportunistic pathogen, is known for its intrinsic resistance to a broad spectrum of antibiotics, including β-lactams, quinolones, aminoglycosides, and polymyxins (7, 8). It has been prioritized by the World Health Organization due to its tendency to develop multidrug resistance (9). As a result of such high levels of resistance and the severity of *P. aeruginosa* infections, phage therapy has emerged as a promising tool in the treatment of these infections, greatly increasing research efforts in characterizing *Pseudomonas* phages (10–13). Among well-studied *Pseudomonas* phages, Pf4 is notable for its seemingly mutualistic relationship with its host. It also plays a role in superinfection exclusion (SIE), targeting the Type IV Pilus (T4P), an essential structure for *P. aeruginosa* twitching motility and associated virulence, for initial infection and subsequent SIE (14–19).

SIE is a mechanism by which prophages inhibit infection of their bacterial host by other similar phages. Prophage genes frequently encode proteins that can interfere with the infection mechanisms of invading phages, often inducing significant alterations to the host’s surface structures (20–24). These surface modifications have far-reaching implications for the fitness and survival of the host bacterium (25). The following study investigates the relationship between Pf4 phage and its *P. aeruginosa* host, exploring the role of antibiotics in the induction of prophages and the consequent fitness effects arising from prophage induction and the activation of SIE mechanisms.

## Results

### Phylogenetic Evidence Suggests Evolutionary Maintenance of Pf Phage Genes

To demonstrate the significance of Pf4 phage as a driver of *P. aeruginosa* evolution and behavior, we quantified the prevalence of integrated Pf prophages in *P. aeruginosa*. We screened 405 *P. aeruginosa* whole genome sequences (WGS) from the Pseudomonas Genome Database for the presence of the major coat protein of Pf phages, *coaB* (26, 27). We found that approximately 53% (212/405) contained a region with at least 90% sequence similarity and complete alignment coverage for the *coaB* gene. Using Roary for core genome alignment, we mapped the presence of *coaB* genes onto a phylogenetic tree (Fig. 1A), which showed significant clustering within phylogenetic groups, indicating a selection for Pf prophages in *P. aeruginosa*.

**Fig. 1.**
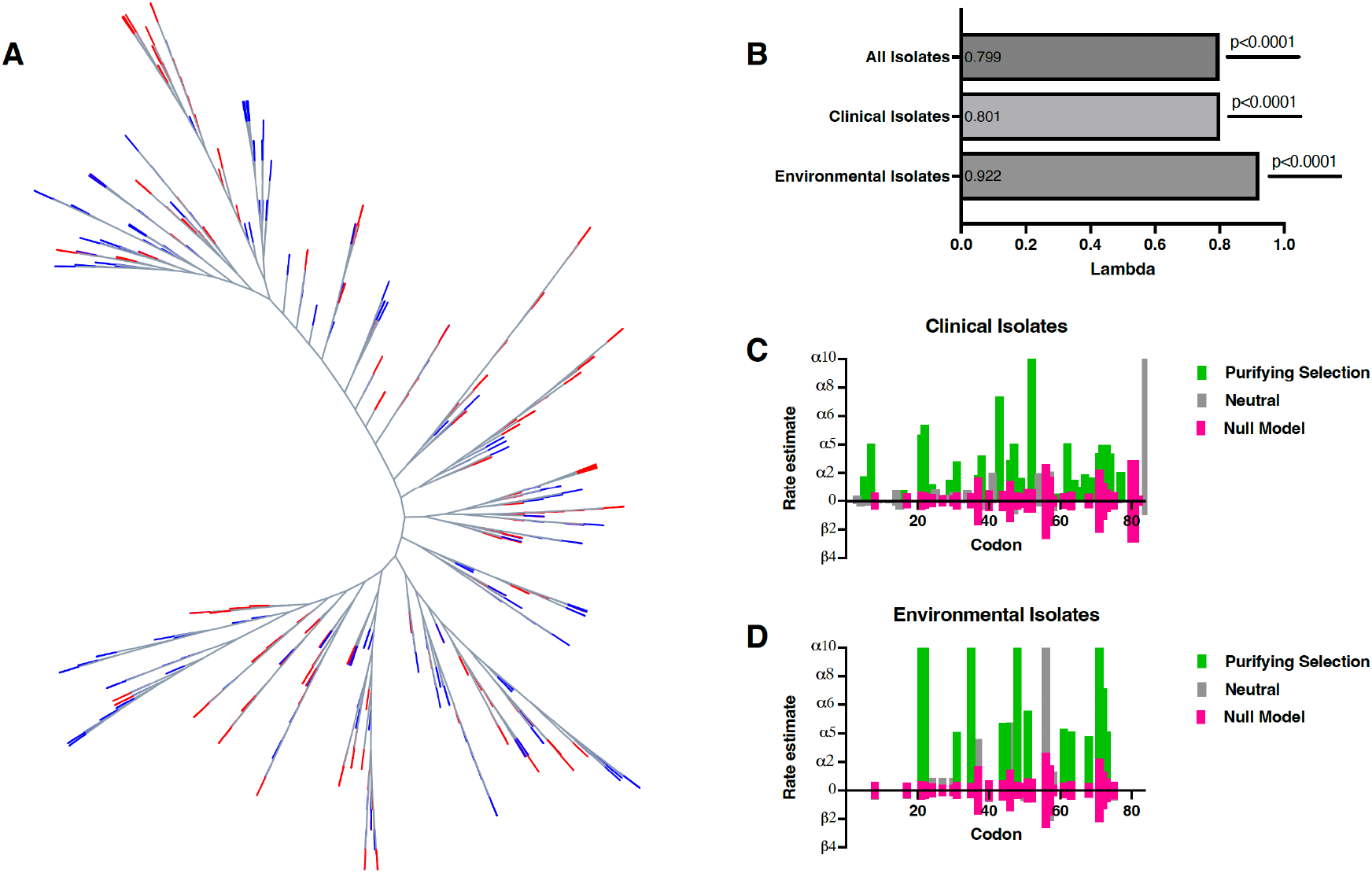
Phylogenetic analysis of *P. aeruginosa* isolates shows a significant relationship with Pf phages. (A) Unrooted phylogenetic tree of n= 405 *P. aeruginosa* isolates, branch colors demonstrate presence (blue) or absence (red) of *coaB* Pf phage gene. (B) Pagel’s lambda values for phylogenetic signal analysis of *coaB* gene presence for n= 405 total isolates, n= 364 clinical isolates, and n=41 environmental isolates. (C-D) Synonymous (alpha) and non-synonymous (beta) mutation rates in *coaB* gene as predicted by fixed effects likelihood model for (C) clinical isolates and (D) environmental isolates of *P. aeruginosa*. Statistical analysis for (B) Pagel’s lambda was performed by likelihood ratio test.

Next, to assess the relationship between *coaB* gene presence and phylogeny, we performed a phylogenetic signal analysis using Pagel’s lambda values. The analysis yielded a Pagel’s lambda value of 0.799 with highly significant p-values (p < 2.2×10^−16^), demonstrating robust phylogenetic signals (Fig. 1B). We categorized these genomes by isolation sources into clinical (n=364) and environmental (n=41) groups, revealing that clinical isolates retained a similar lambda value (0.801), while environmental isolates showed a higher lambda value (0.922). These results suggest the evolutionary conservation of Pf prophages in environmental isolates, supporting their important role in SIE and host fitness.

Examination of selection pressures on the *coaB* gene using both fixed effects likelihood and mixed effects models of evolution showed purifying selection at approximately 50% of the non-invariant sites. A singular site in clinical isolates displayed episodic positive selection. Clinical isolates had a marginally higher percentage of sites under purifying selection (53%) compared to environmental isolates (50%), but environmental isolates exhibited higher rates of purifying selection per site (Fig. 1C-D), indicating stronger selective pressure in these isolates.

### Sub-lethal Concentrations of Antibiotics Induce Prophage via the Activation of DNA SOS Response

To identify sublethal antibiotic concentrations for *P. aeruginosa* consistent with our study’s objectives, we conducted liquid broth dilution assays (Fig. S1). We determined the minimum inhibitory concentration (MIC) and MIC_20_ values for all tested antibiotics (Table S1). MIC_20_ is defined as the minimum antibiotic concentration that inhibits 20% or less of bacterial growth over 16 hours, allowing us to investigate the fitness effects imposed by prophage induction and subsequent activation of SIE while minimizing significant antibiotic-mediated toxicity.

**Fig. S1.**
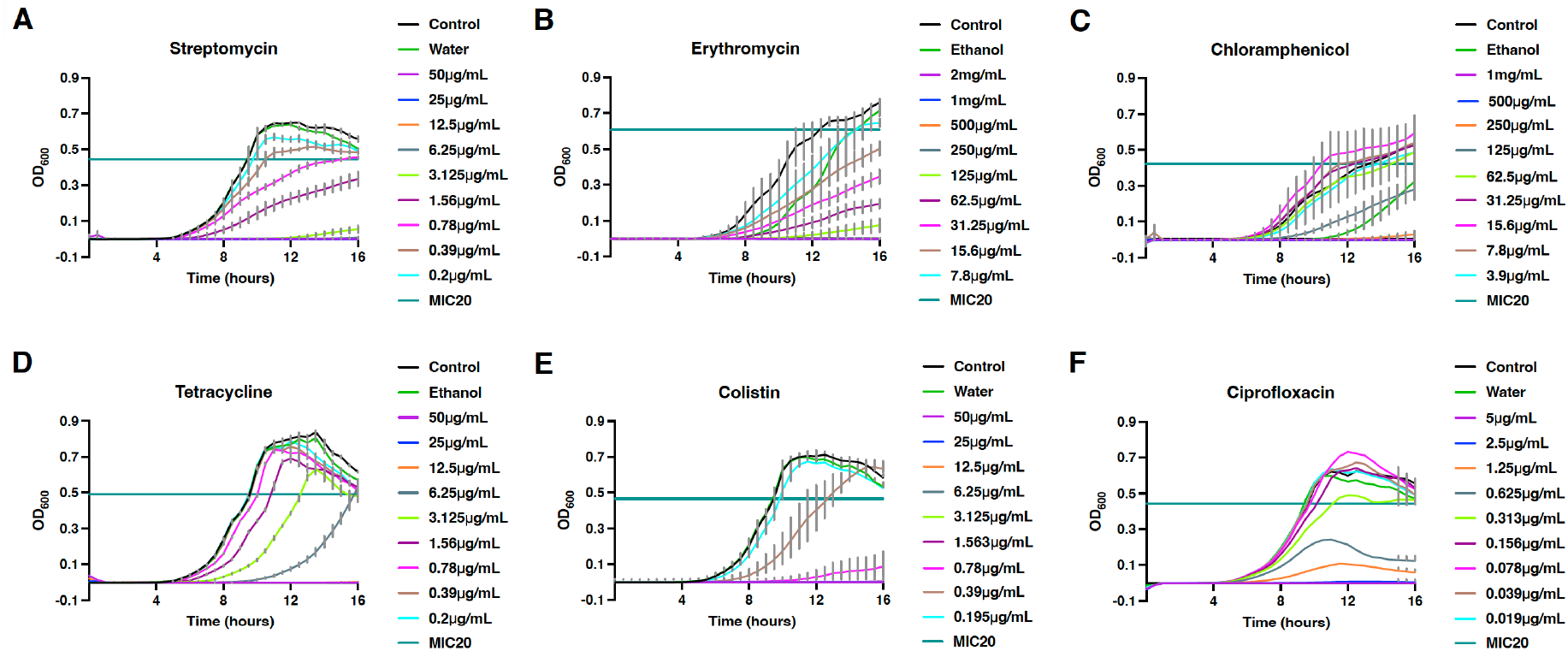
Minimum inhibitory concentrations and MIC_20_ values of various antibiotics for *P. aeruginosa* PAO1 were identified. (A-F) The minimum inhibitory concentration (MIC) for various antibiotics, including (A) Streptomycin, (B) Erythromycin, (C) Chloramphenicol, (D) Tetracycline, (E) Colistin, and (F) Ciprofloxacin, for *P. aeruginosa* PAO1 was identified via liquid broth dilution assay in LB medium. MIC_20_ was calculated as the minimum concentration that inhibited at least 20% of bacterial growth after 16 hours. All graphs (A-F) are representative of n = 3 independent experiments and show the mean with standard error of the mean (SEM) of n = 24 replicates per group.

**Table S1.** Minimum inhibitory concentrations and optimal inducing concentrations of tested antibiotics against *P. aeruginosa* PAO1.

| Antimicrobial Compound | MIC (μg/mL) | MIC<sub>20</sub> (μg/mL) | Optimal Inducing Concentration (μg/mL) |
| --- | --- | --- | --- |
| Streptomycin | 6.25 | 0.78 | 0.39 |
| Erythromycin | 250 | 7.8 | 1.95 |
| Chloramphenicol | 500 | 62.5 | 15.6 |
| Tetracycline | 12.5 | 3.13 | 0.39 |
| Colistin | 1.56 | 0.39 | 0.039 |
| Ciprofloxacin | 2.5 | 0.31 | 0.078 |

After determining MIC_20_ values, we examined the ability of these antibiotics to induce Pf4 phage in *P. aeruginosa* PAO1 at concentrations above and below the identified MIC values. We found that sublethal concentrations of tetracycline, colistin, and ciprofloxacin significantly increased prophage induction compared to spontaneous induction controls (Fig. 2D-F). We also showed that tetracycline lost the ability to induce prophages when in the absence of light (Fig. S2). In contrast, erythromycin, streptomycin, and chloramphenicol did not show a significant increase in Pf4 induction (Fig. 2A-C).

**Fig. 2.**
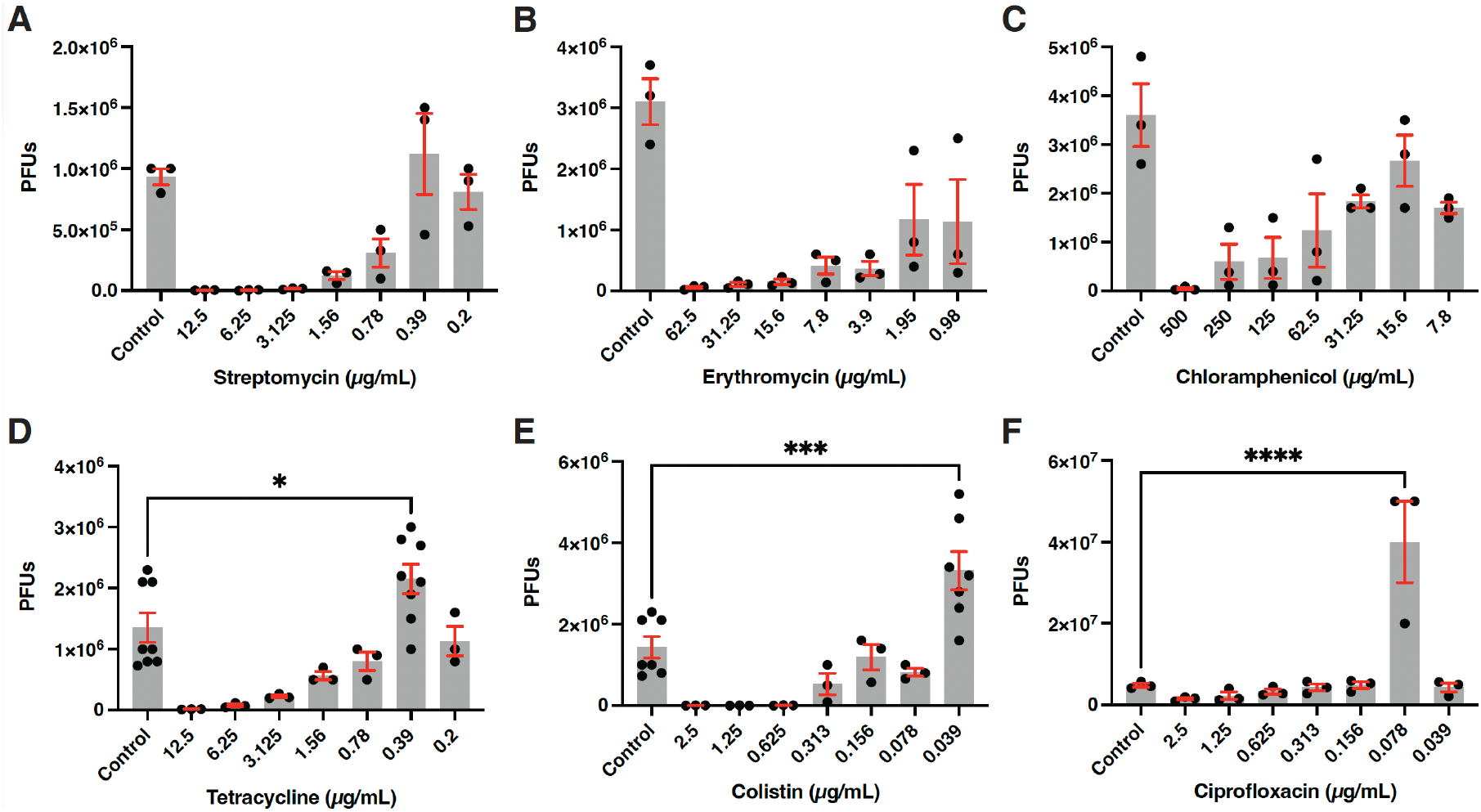
Sub-lethal concentrations of several antibiotics induce Pf4 phage at significantly increased levels. (A-F) Levels of Pf4 induction quantified by plaque assay plated on PAO1ΔPf4 after 4 hours of exposure to a range of antibiotic concentrations for (A) Streptomycin, (B) Erythromycin, (C) Chloramphenicol, (D) Tetracycline, (E) Colistin, and (F) Ciprofloxacin. All graphs (A-F) are representative of n *≥* 3 independent experiments and show the mean with SEM of n *≥* 3 replicates. Statistical comparisons of (A-F) were done via one-way analysis of variance (ANOVA) followed by Dunnett’s multiple comparisons test.

**Fig. S2.**
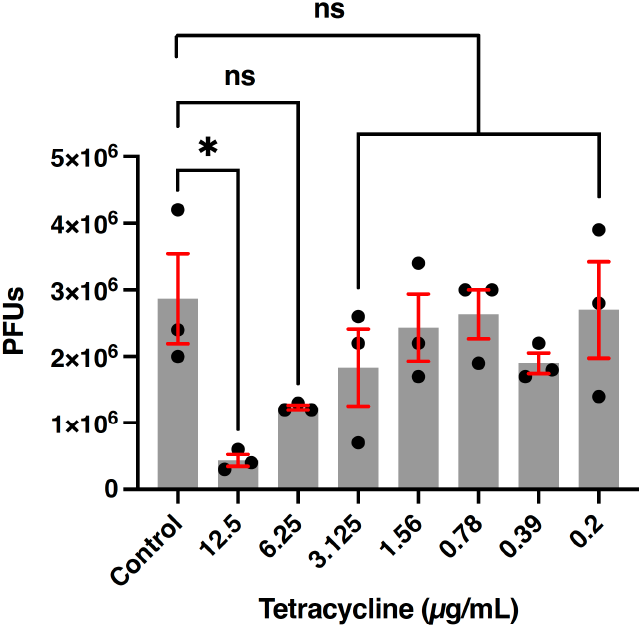
Tetracycline fails to increase Pf4 prophage induction in the absence of light. Pf4 prophage induction levels were measured by plaque assay on *P. aeruginosa* PAO1ΔPf4 when exposed to a range of concentrations of tetracycline and in the absence of light. The graph is representative of n = 3 independent experiments and shows the mean ± SEM of n = 24 replicates per group. Statistical comparisons were performed via a one-way ANOVA followed by Dunnett’s multiple comparisons test.

To understand the mechanisms of Pf4 phage induction by these antibiotics, we used PAO1 transposon mutant strains lacking functional *recA* or *lexA* genes, which represent a key activator and a repressor of the DNA SOS response, respectively. In the background of these mutants, the inducing antibiotics lost their ability to significantly induce prophages above control levels (Fig. 3A-C). To further understand the impact of these antibiotics on bacteria at non-effective concentrations, we assess the production of reactive oxygen species (ROS) in liquid cultures in the presence of antibiotics at their optimal inducing concentrations (Table S1) by measuring 2’7’-dichlorofluorescin (DCF) fluorescent signal intensity. We showed that tetracycline, colistin, and ciprofloxacin induced ROS above control levels (Fig. 3D).

**Fig. 3.**
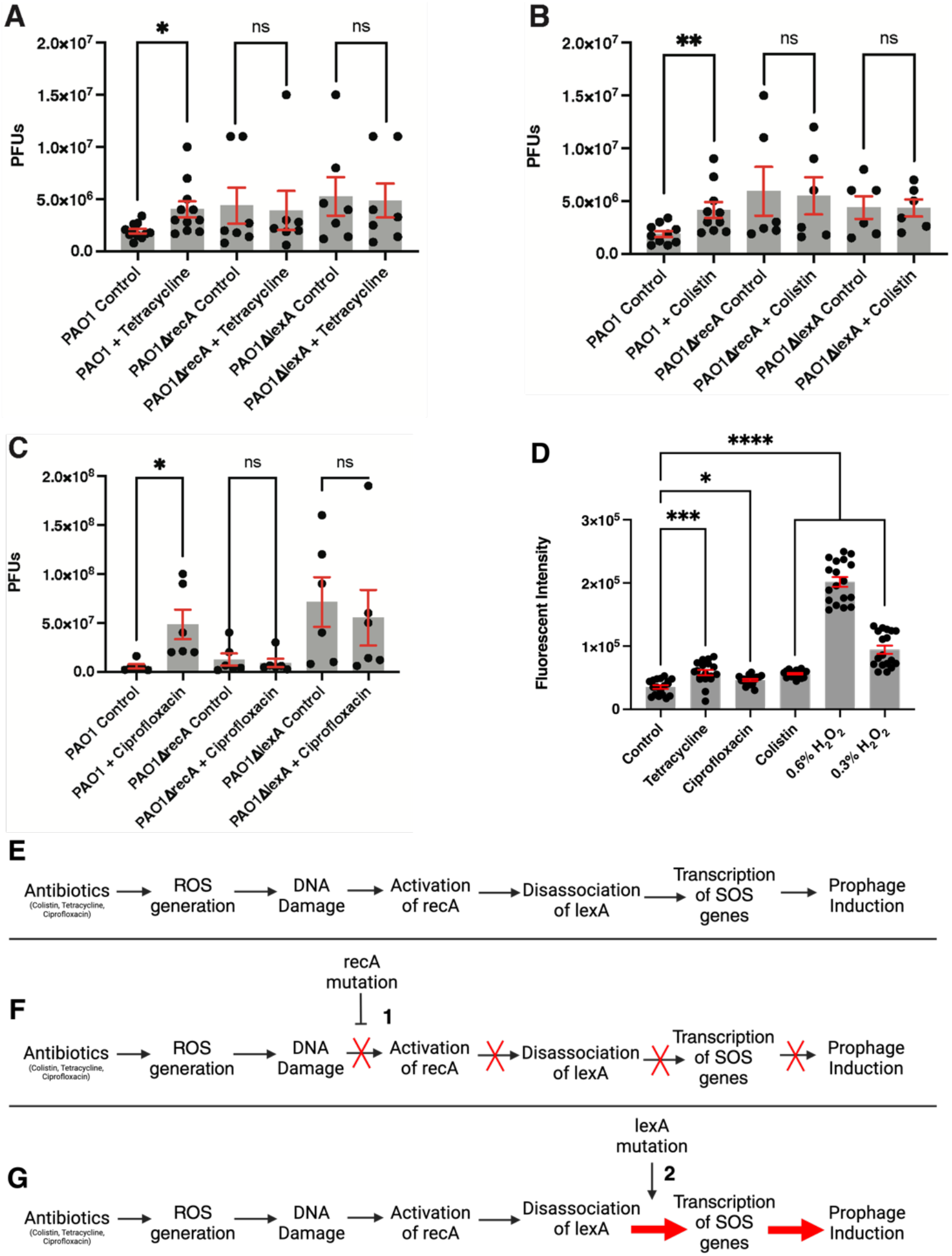
Induction of Pf4 phage by sublethal concentrations of antibiotics involves the generation of ROS and is dependent on bacterial SOS response. (A-C) The ability of antibiotics to induce Pf4 phage when lacking key activators or repressors of the DNA SOS response was tested by plaque assay using PAO1ΔPf4 as the host strain in the presence of (A) Tetracycline, (B) Colistin, or (C) Ciprofloxacin. (D) Reactive oxygen species were quantified via DCF fluorescence after exposure to various antibiotics at their optimal inducing concentrations, as stated in Table S1, for 15 minutes and exposed to ambient light. (E-F) Hypothetical mechanisms of antibiotic-mediated induction of prophages in wildtype PAO1 (A), PAO1ΔrecA, where a mutation resulting in loss of functional RecA causes inhibition (1) of all RecA dependent steps (F), and PAO1Δ*lexA*, where mutation resulting in loss of functional LexA causes the constitutive expression (2) of SOS genes, and increased prophage induction masking the antibiotic mediated inducing effect (G). Statistical comparisons were performed via Welch’s ANOVA followed by Dunnett’s multiple comparisons test. All graphs (A-D) are representative of n *≥* 3 independent experiments and show the mean with SEM of n *≥* 3 replicates. Statistical comparisons of (A-C) were done by two-tailed Student’s t-test. Statistical comparisons of (G) were done via Welch’s ANOVA, followed by Dunnett’s multiple comparisons test.

### Pf4 Superinfection Influences Host Fitness

After identifying conditions that induce Pf4 phage, we evaluated the impact of prophage induction on bacterial fitness. Pf4 superinfection did not significantly affect the growth rate of *P. aeruginosa* PAO1 in enriched (Fig. 4A) or minimal media (Fig. 4B) across a range of phage concentrations. However, superinfected *P. aeruginosa* exhibited significantly reduced twitching motility compared to both uninfected PAO1 and PAO1ΔPf4 (Fig. 4C). By assessing biofilm formation through crystal violet staining, we found that superinfecting *P. aeruginosa* PAO1 with a high multiplicity of infection (MOI) of 1, 5, or 10 of Pf4 did not significantly alter biofilm adherence (Fig. 4D-E). In contrast, low MOIs (0.01, 0.1) significantly increased biofilm adherence. Additionally, PAO1ΔPf4 and PAO1ΔPilC showed significantly impaired biofilm adherence compared to the wildtype after 24 hours.

**Fig. 4.**
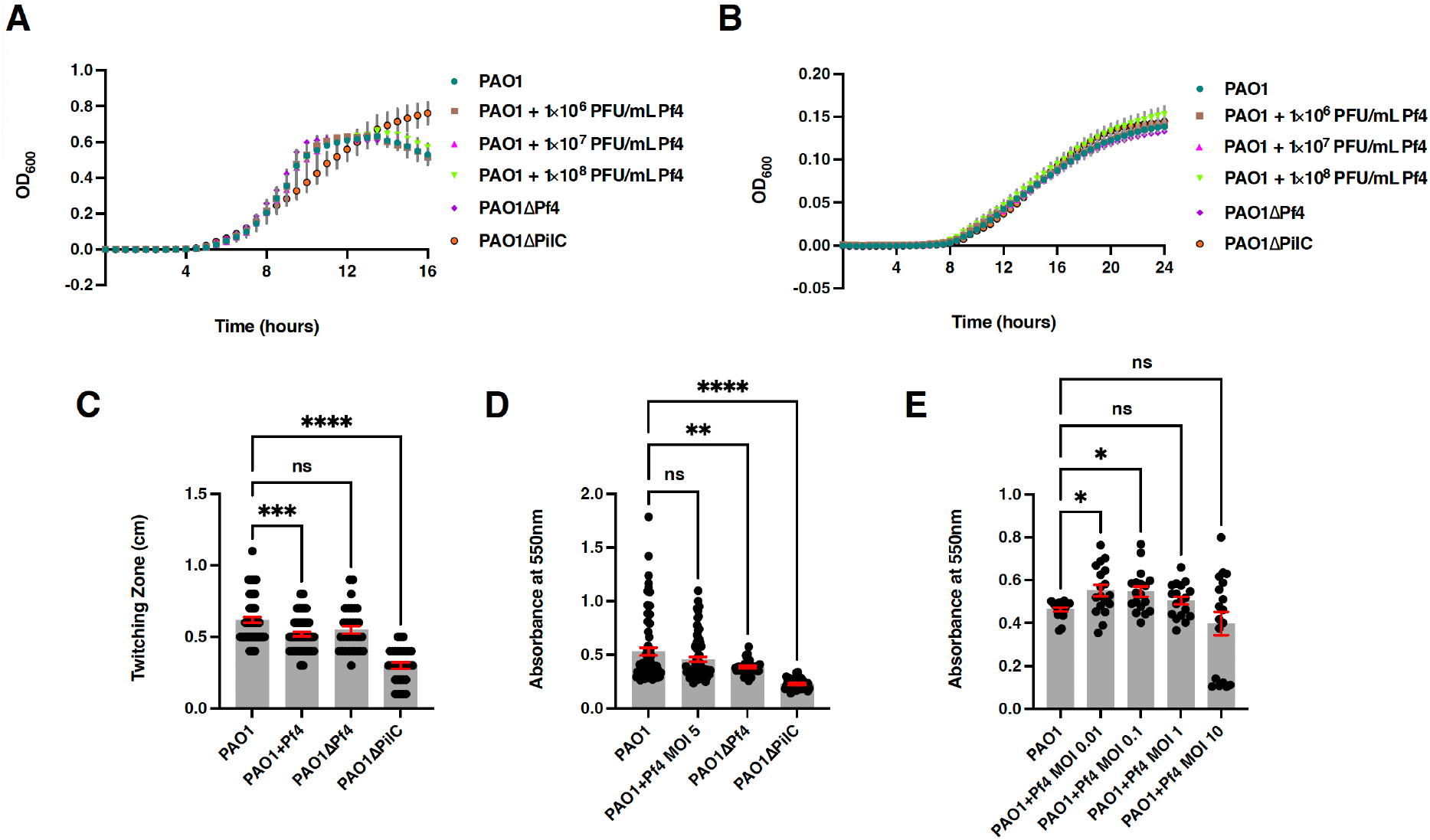
Pf4 superinfection significantly affects the fitness of *P. aeruginosa* host. (A-B) Growth curves of *P. aeruginosa* PAO1 untreated or superinfected with a range of concentrations of Pf4 phage as well as *P. aeruginosa* mutant strains PAO1ΔPf4 and PAO1ΔPilC in (A) enriched medium and (B) minimal medium. (C) Twitching motility assay of *P. aeruginosa* PAO1 untreated or superinfected with Pf4 phage at MOI 10 and mutants, PAO1ΔPf4 and PAO1ΔPilC. (D-E) Biofilm adhesion of *P. aeruginosa* PAO1 untreated or superinfected with Pf4 at MOIs ranging from 0.01-10, and mutant strains, measured by crystal violet staining. All graphs (A-E) are representative of n *≥* 3 independent experiments and show the mean ± SEM of n *≥* 3 replicates. Statistical comparisons of (C-E) were done via one-way ANOVA followed by Dunnett’s multiple comparisons test.

To determine the impact of Pf4 superinfection on the ability of *P. aeruginosa* to evade phagocytosis, macrophage-mediated uptake was quantified employing a gentamicin protection assay followed by cell lysis and enumeration of intracellular bacteria by plating for colony-forming units (CFUs). The results revealed a significant 7-fold increase in macrophage uptake of *P. aeruginosa* PAO1 cells that were previously superinfected with Pf4 at MOI 10 compared to uninfected controls (Fig. 5A). Similarly, PAO1ΔPilC showed a nearly 8-fold increase in uptake. When testing the effect of superinfection by a Pf4 phage isolated from a clinical isolate, *P. aeruginosa* 264, there was no significant increase in phagocytosis compared to the uninfected host. Furthermore, we investigated the effect of Pf4 superinfection on *P. aeruginosa* clearance from macrophages by plating CFUs from lysed macrophages at 2, 4, 6, and 24 hours post-infection. Our results revealed that both superinfected PAO1 and PAO1ΔPilC were cleared at a faster rate than uninfected PAO1 (Fig. 5B). Furthermore, we determined that infecting PAO1 with Pf4 at MOI 10 at the time of infection resulted in a significant decrease in phagocytosis (Fig. 5C).

**Fig. 5.**
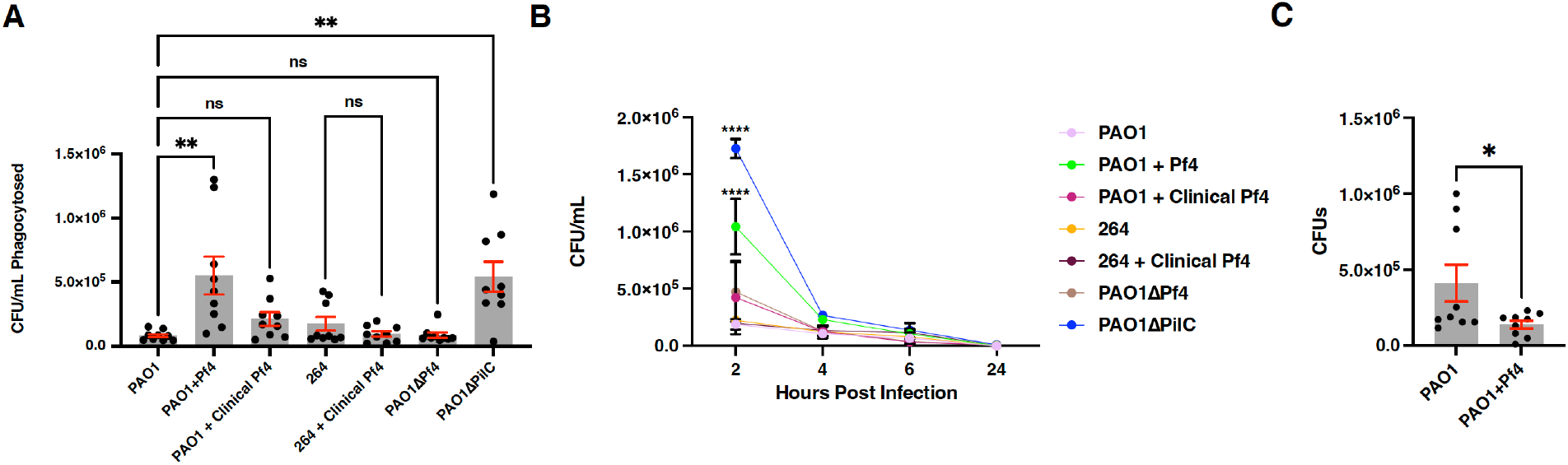
Macrophage-mediated phagocytosis and clearance of *P. aeruginosa*. (A) Phagocytosis and (B) clearance of *P. aeruginosa* PAO1 untreated or previously superinfected by Pf4 or clinical Pf4 at MOI 10 for 18 hours, along with mutant strains and clinical isolate *P. aeruginosa* 264, tested by gentamicin protection assay followed by plating colony forming units. (C) Macrophage Phagocytosis of PAO1 untreated or superinfected with Pf4 at MOI 10 at the time of infection. All graphs (A-C) are representative of n *≥* 3 independent experiments and show the mean ± SEM of n *≥* 3 replicates. Statistical comparison (A) was done via one-way ANOVA followed by Dunnett’s multiple comparisons test, except the comparison between 264 and 264 + clinical Pf4 in (A), which was completed by a two-tailed Student’s t-test. Comparisons in (B) were performed via two-way ANOVA followed by Dunnett’s multiple comparisons test. Paired Student’s t-test was performed in (C).

## Discussion

The findings of this study demonstrate the complexity of phage-host interactions, wherein the induction of prophages by antibiotic-producing microbes can trigger phenotypic changes in the bacterial host that have broader implications for the rest of the population. They also suggest a valid evolutionary mechanism that may explain the production of sub-lethal concentrations of antibiotics by competing bacterial species in the environment by demonstrating that these concentrations can induce prophages in *P. aeruginosa* and lead to alterations in fitness (Figure 6).

**Fig. 6.**
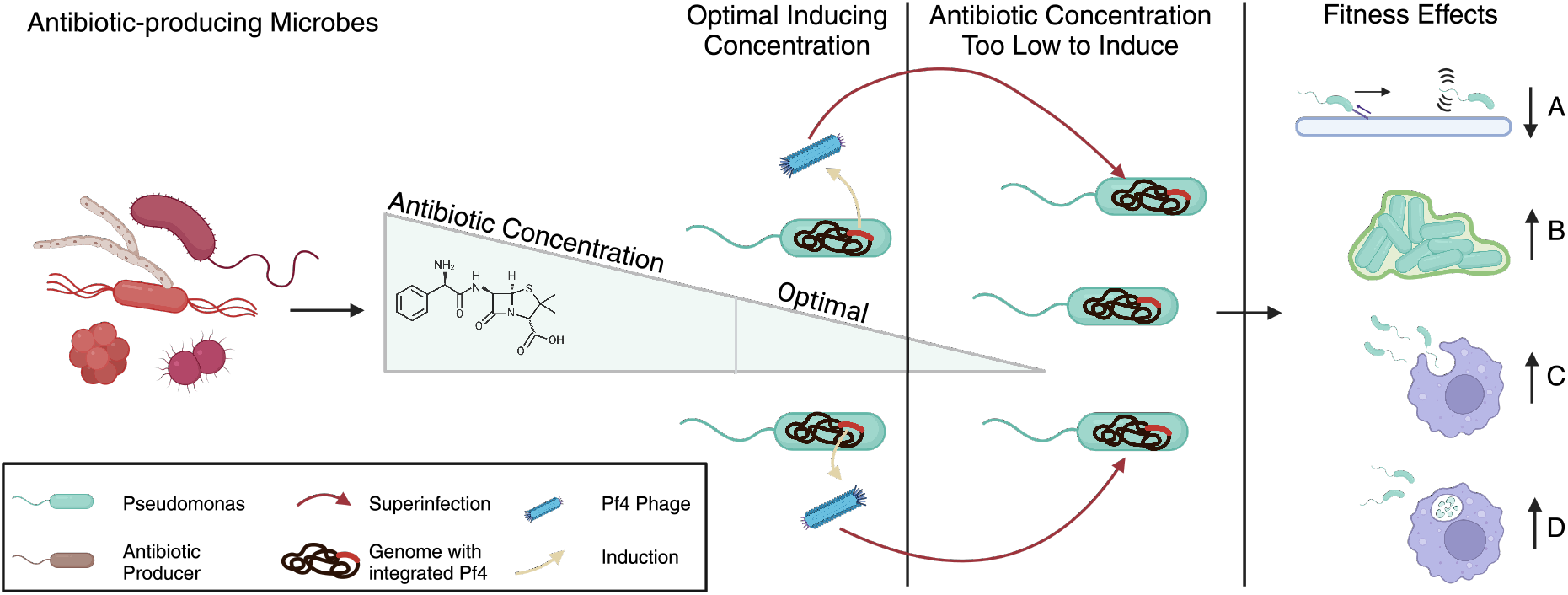
Proposed model for the evolutionary role of production of sublethal concentrations of antibiotics by antibiotic-producing microbes in the environment. Antibiotic producers release antibiotics, which interact with neighboring bacterial populations. Some members of these populations may be located at a distance that exposed them to an optimal prophage-inducing concentration, leading to prophage induction. These induced prophages will then go on to attempt to superinfect the rest of the population, resulting in significant fitness effects. These effects, which are dependent on the environment and the concentration of induced phages, include (A) loss of twitching motility, (B) increase in biofilm adherence, (C) increase in phagocytosis, and (D) increase in clearance by macrophages.

The genomic analyses conducted in this study provided phylogenetic evidence of evolutionary maintenance for Pf prophage regions (Fig. 1). This concept has previously been hypothesized due to the evolutionary benefits that Pf phages, namely Pf4 phages, confer to their host through improvements in biofilm stability (27–29). Additionally, our study showed that Pf phage presence was more strongly correlated with phylogeny in environmental isolates than in clinical isolates (Fig. 1B). We believe this observation may be attributed to the capacity for Pf4 superinfection to alter *P. aeruginosa* macrophage evasion and clearance, a fitness consequence that differentially impacts clinical isolates and environmental isolates. The dominance of purifying selective pressures on these genes indicates the functional importance of these genes not only for maintenance of the Pf phage but also for bacterial fitness, suggesting that bacterial evolution and viral evolution likely together support the maintenance of Pf phages in *P. aeruginosa*.

Next, we found that sub-therapeutic concentrations of several antibiotics, including tetracycline, colistin, and ciprofloxacin, can induce Pf4 phage at significantly higher levels than what occurs spontaneously (Fig. 2). In support of the results obtained in our study, ciprofloxacin, a synthetic antimicrobial, has previously been shown to induce prophages via activation of the SOS response (30–32). To the best of our knowledge, this study is the first report of tetracycline and colistin, both naturally produced antibiotics, acting as inducers of prophages.

Existing literature suggests that ROS can act as potent inducers of prophages (33). Additional evidence supports the induction of ROS by tetracycline and colistin (34–38). Therefore, we explored the possibility that ROS generation and subsequent activation of bacterial SOS response could explain prophage induction by these antibiotics. First, tetracycline lost the ability to induce Pf4 in the absence of light (Fig. S2), aligning with previous reports that tetracycline-mediated ROS generation is dependent on light activation (36–38). Next, we showed that ROS generation by these antibiotics occurred at their optimal inducing concentrations (Fig. 3D). Furthermore, by showing that the inducing antibiotics lost the ability to increase prophage induction in mutants lacking an essential activator (recA) or suppressor (lexA) of the DNA SOS response pathway compared to untreated controls, our study demonstrated that all inducing antibiotics rely on activation of the DNA SOS response to successfully increase prophage induction (Fig. 3A-C). This finding again confirms that ciprofloxacin induces prophages via activation of the DNA SOS response. Collectively, our results provide a potential mechanism by which tetracycline and colistin induce prophages. Both antibiotics induce ROS, which contribute to DNA damage and, in turn, activate the DNA SOS response, ultimately leading to the induction of Pf4 phages.

Having established that various antibiotics can induce the production of Pf4 phage through a validated mechanism, we next investigated how Pf4 phage superinfection affects the fitness of *P. aeruginosa*. We showed that superinfection by Pf4 phage, *P. aeruginosa*, exhibited a significant reduction in twitching motility (Fig. 4C), aligning with previous reports (20, 29). Further, superinfecting *P. aeruginosa* with high concentrations of Pf4 phage (1 × 10^8^ PFU/mL) had no significant effect on biofilm adhesion after 24 hours (Fig. 4D). This finding presents a contrast to other research, which has reported improved biofilm adhesion in PAO1 when supplemented with a lower concentration of Pf4 phage (1 × 10^6^ PFU/mL) (39). These contrasting results led us to hypothesize that Pf4 phage’s effect on PAO1 biofilm adhesion is dependent on the phage concentration. We confirmed this by demonstrating that at low MOIs, superinfection by Pf4 increased biofilm formation significantly (Fig. 4E), aligning with the previous report. However, at higher MOIs, Pf4 phage is unable to significantly improve biofilm formation.

Our study further suggests that the concentration-dependent effect on biofilm adherence may be the result of two competing influences. First, previous studies have demonstrated that Pf4 phages, when secreted by *P. aeruginosa*, may act as a scaffold for biofilms, thereby bolstering adhesion (39). Such a mechanism is supported by our results demonstrating that *P. aeruginosa* PAO1ΔPf4 had a significant decrease in biofilm formation compared to *P. aeruginosa* PAO1. Conversely, the exclusion of superinfecting Pf4 phage has been linked to specific interactions with the PilC protein, an essential protein for the assembly and stability of the T4P, which is important for biofilm adherence (17). Consistent with these previous reports, a PAO1 strain carrying a PilC knockout mutation used in our study exhibited a significant decrease in biofilm formation.

Thus, at lower concentrations, Pf4 phage may enhance biofilm formation by providing additional stability to biofilms, whereas at higher concentrations, the inhibitory effects on the T4P assembly and stability induced by SIE of Pf4 become more dominant, eliminating any improvements Pf4 phage may provide to biofilm adherence.

Next, Pf4 phage superinfection resulted in enhanced macrophage-mediated bacterial uptake and clearance (Fig. 5A-B). The increase in phagocytosis could be explained by two possible mechanisms: either superinfection by Pf4 phage undermines the ability of *P. aeruginosa* to evade macrophages as a result of the loss of its T4P, suggesting that the T4P might play an important role in immune evasion, or superinfection by Pf4 phage alters the bacterial surface structure in some way that improves its recognition by macrophages. Previous studies have suggested that Pf4 phages may inhibit the phagocytosis of *P. aeruginosa* by macrophages when cells were treated with Pf4 and *P. aeruginosa* PAO1 simultaneously (40, 41), a result we were able to confirm in this study (Fig. 5C). However, the effects of Pf4 phage superinfection on *P. aeruginosa* evasion of phagocytosis when *P. aeruginosa* had been pre-infected with Pf4 phage, allowing superinfection exclusion mechanisms to activate, had not been examined prior to this study. The mechanism by which the superinfected phenotype exhibits enhanced clearance remains to be investigated.

Furthermore, when testing the capacity for Pf4 phages isolated from clinical *P. aeruginosa* isolates to enhance phagocytosis and clearance, we saw no significant effect. It is possible that clinical isolates have evolved to mitigate these fitness costs incurred by Pf4 phage as a result of their exposure to unique selection pressures. These adaptations could reasonably explain the evidence of episodic positive selection, found only in clinical isolates, as exposure to strong selection pressures such as antibiotics or the immune system would provide the basis for positive selection.

Taken together, the results of our study also suggest an explanation for the synergistic effects observed between antibiotics and lytic phages during phage therapy. We have demonstrated that prophage induction can lead to fitness defects in bacterial communities due to the activation of SIE mechanisms. By using antibiotics, even at sub-therapeutic concentrations, these mechanisms might be stimulated as prophages are induced, potentially suppressing the emergence of phage-resistant mutants and increasing the effectiveness of phage therapy. This complex interplay might enhance therapeutic outcomes beyond the expected additive inhibitory effects of antibiotics and lytic phage therapy.

Furthermore, the implication of our results extends beyond the environmental and clinical settings and can affect other circumstances where bacteria are exposed to low concentrations of antibiotics. For example, low concentrations of antimicrobials are frequent pollutants found in aquatic ecosystems, ultimately affecting marine life and making their way to urban water systems (42, 43). The aquatic environment is also a hotspot for bacterial communities, which, when exposed to sub-inhibitory concentrations of antibiotics, may activate prophages, leading to SIE mechanisms that reduce bacterial fitness and impact microbial community dynamics. Such a scenario could result in altered microbial interactions and potential shifts in ecosystem functions, emphasizing the broader ecological impact of antibiotic pollution other than the development of antimicrobial resistance or a direct antimicrobial effect. A similar scenario of prophage-mediated fitness cost among bacteria may be transiently observed in humans and animals at the end of the antibiotic treatment when its levels fall below the therapeutic dose.

In summary, our findings provide insights into the complex role of Pf4 phage in *P. aeruginosa* physiology and evolution. The evidence of strong phylogenetic signal of Pf4 prophage genes within *P. aeruginosa* genomes suggests a potential symbiotic contribution to the host, especially in environmental isolates and when phages are present in lower concentrations. The findings that antibiotics like tetracycline, colistin, and ciprofloxacin can induce prophages at levels surpassing spontaneous induction serve as a potential evolutionary explanation for the production of sublethal concentrations of antibiotics in the environment, which may affect not only bacterial populations but, as previously discovered, fungi can also be affected by this mechanism (44, 45). Lastly, the study uncovers a highly complex interaction between Pf4 superinfection and *P. aeruginosa* fitness, affecting numerous phenotypes and doing so in a concentration, spatial, and temporally dependent manner. Taken together, these findings suggest the importance of further research into the ecological and clinical significance of phage-host dynamics.

## Materials and Methods

### Bacterial/Phage Stocks and Culturing

All bacterial strains are listed in Table S2 and were cultured in Lennox broth (LB) (10 g of LB powder (Apex Bioresearch Products) dissolved in 1L of nanopure H_2_O) at 37°C and shaking at 220 revolutions per minute (RPM) overnight. Pf4 phage stocks were prepared via prophage induction, as outlined below. Phage stocks were concentrated through three successive incubations with PAO1ΔPf4 permissive host and centrifuged at 22,000 × g for 60 minutes. Phage stocks were stored at 4°C in SM buffer (8 mM MgSO_4_·7H_2_O, 100 mM NaCl, 50 mM Tris-HCl pH 7.4, dissolved in 1 L of nanopure H_2_O). RAW 264.7, immortalized macrophages (ATCC TIB-71) were maintained utilizing Dulbecco’s Modified Eagles Medium (DMEM) (GenClone) media supplemented with 10% fetal bovine serum (FBS) (GenClone) and 1% Penicillin-Streptomycin (GenClone) at 37°C and 5% CO_2_.

**Table S2.** Bacterial strains used in this study.

| Strain | Description | Source |
| --- | --- | --- |
| *Pseudomonas aeruginosa* PAO1 | Reference Strain | ATCC |
| *P. aeruginosa* PAO1ΔPf4 | *P. aeruginosa* PAO1 strain with deletion of Pf4 phage | Secor Lab, (20) |
| *P. aeruginosa* PAO1ΔpilC | *P. aeruginosa* PAO1 strain with transposon mutation interrupting pilC gene | Manoil Library, (46) |
| *P. aeruginosa* PAO1ΔrecA | *P. aeruginosa* PAO1 strain with transposon mutation interrupting recA gene | Manoil Library, (46) |
| *P. aeruginosa* PAO1ΔlexA | *P. aeruginosa* PAO1 strain with transposon mutation interrupting lexA gene | Manoil Library, (46) |
| *P. aeruginosa* 264 | Multidrug-resistant clinical isolate of *P. aeruginosa* | CDC ARIsolate Bank, (47) |

### Phage Isolation/Plaque Assay

Pf4 phage was isolated from *P. aeruginosa* PAO1. Bacteria were grown overnight, diluted to 1×10^7^ CFU/mL, and treated with 0.5 µg/mL of Mitomycin C (Fisher Scientific) for 4 hours at 37°C. Following the incubation, the sample was centrifuged at 10,000 × g to pellet bacteria. The supernatant was transferred to a new tube and filtered using 0.22 µm syringe filters. The presence of phage was confirmed using plaque assay as described previously by plating on *P. aeruginosa* PAO1ΔPf4 (48). The plates were incubated at 37°C for 18 hours.

### Phage Quantification

After a purified lysate was produced, phage concentration was determined using a plaque titer assay. Phage lysates were serially diluted. A 10 µL aliquot of each dilution was mixed with 3 mL of melted LB agar that has cooled to 55°C and 1 mL of *P. aeruginosa* PAO1ΔPf4, followed by spreading on LB plates forming a top agar layer. Plates were incubated at 37°C for 18 hours. Following incubation, plaques were counted, and titer was calculated.

### Antibiotic MIC Assays

The minimum inhibitory concentration (MIC) in LB broth of several antibiotics was identified for *P. aeruginosa* PAO1 as previously described (49). To make stock concentrations, Colistin sulfate (Acros organics), Streptomycin sulfate (Thermo Fisher Scientific), and Ciprofloxacin (Tokyo Chemical Industry) were dissolved in H_2_O. Tetracycline hydrochloride (Fisher Scientific) was resuspended in 70% ethanol (Fisher Bioreagents). Chloramphenicol (Sigma Aldrich) and Erythromycin (Sigma Aldrich) were resuspended in 100% ethanol (Fisher Bioreagents). Antibiotics were then diluted from stock to the relevant starting concentrations in LB broth and were then diluted 1:2 across a 96-well plate. An overnight culture of *P. aeruginosa* PAO1 was set to an Optical Density (OD_600_) of 1.0 (2 × 10^8^ CFU/mL) and then diluted in LB 1:500. A 50 µL aliquot of this dilution was added to each well, bringing the total volume to 100 µL and diluting the antibiotic concentration by 1:2. The OD_600_ readings were monitored in a Tecan M200Pro microplate reader for 16 hours at 37°C with 5 seconds of orbital shaking and readings taken every 30 minutes. The MIC was calculated as the lowest concentration that prevented the growth of *P. aeruginosa* PAO1 over the course of treatment. MIC_20_ was calculated as the lowest concentration that inhibited bacterial growth by 20% at the end of 16 hours.

### Prophage Induction Assays

#### General Procedure

The capacity for a given antibiotic to induce Pf4 was evaluated using a modified plaque assay. An overnight culture of *P. aeruginosa* PAO1 was diluted to OD_600_ = 0.05 (1×10^7^ CFU/mL) and treated with a range of antibiotic concentrations or left untreated to serve as a control for spontaneous prophage induction. The bacteria were then incubated at 37°C for 4 hours shaking at 220 RPM. Following the incubation, samples were immediately centrifuged at 13,500 × g for 5 minutes to pellet bacteria. The supernatant was carefully removed and aliquoted into new tubes, followed by vortexing and serial dilution. A 10 µL aliquot of each dilution was spotted on LB plates with *P. aeruginosa* PAO1ΔPf4 using the above-described top agar layer method. The plates were incubated at 37°C for 18 hours followed by plaque enumeration as described (48).

### Induction with SOS Response Mutants

The capacity for all inducing antibiotics to induce Pf4 phage in the absence of functional recA and lexA proteins was evaluated as described in the general procedure with slight modifications. Overnight cultures of *P. aeruginosa* PAO1, PAO1ΔrecA, and PAO1ΔlexA were treated with only the optimal inducing concentrations as determined in this study or left untreated to serve as a control. All other steps were carried out as described in the general procedure.

### ROS Detection

Induction of reactive oxygen species (ROS) in bacteria by prophage-inducing antibiotics was evaluated by measuring DCF fluorescent signal intensity as done previously (50). Briefly, an overnight culture of *P. aeruginosa* PAO1 was set to OD_600_ = 1 (2×10^8^ CFU/mL) in 15 mL. The bacterial suspension was then centrifuged at 4350 × g for 30 minutes to pellet bacteria. The supernatant was removed, and the pellet was washed twice with sterile PBS (Quality Biological) to remove any remaining media. The pellet was then resuspended in 15 mL of PBS. 30 µL of freshly made 100mM 2’,7’-Dichlorofluorescin Diacetate (DCFH) (MilliporeSigma) resuspended in Dimethyl sulfoxide (DMSO) (Fisher Bioreagents) was added to the suspension for a final concentration of 20 µM. The tube was covered in foil to avoid light exposure and incubated with shaking at 220 RPM for 40 minutes. The bacterial suspension with DCFH was then split into nine 1 mL aliquots and ROS-inducing conditions were added to the suspension. An untreated control received only sterile PBS, and 0.6% H_2_O_2_ and 0.3% H_2_O_2_ were used as a positive control. Samples were immediately incubated at 37°C with light exposure for 15 minutes. Fluorescent intensity was then quantified by a Tecan M200Pro microplate reader reading from the top with the gain set to 70. Absorbance at OD_600_ was also recorded and was used to normalize fluorescent intensity to cell density.

### Twitching Motility Assay

Twitching motility was assessed using a well-established protocol (51). Overnight cultures of *P. aeruginosa* PAO1, PAO1ΔPf4, and PAO1ΔPilC were diluted to a concentration of 1×10^6^ CFU/mL in 10 mL of LB broth. One 10 mL aliquot of *P. aeruginosa* PAO1 was then infected with 100 µL of 1×10^9^ PFU/mL of Pf4 phage (Final concentration of 1×10^7^ PFU/mL). The samples were then incubated at 37°C, shaking at 220 RPM for 18 hours. Following the incubation, the samples were stabbed through 1.5% LB agar plates until in contact with the plastic using a sterile pipette tip. These plates were then incubated at 37°C for 18 hours to allow twitching motility along the plastic interface. Next, the agar was carefully removed from the plate using a spatula, and the twitching zones were stained with 0.1% crystal violet (Fisher Scientific) for 15 minutes. Excess stain was washed away with deionized water, and twitching zones were measured using a ruler and expressed in millimeters.

### Biofilm Adherence Assay

Biofilm adherence was assessed using a crystal violet staining assay as described previously (39). Overnight cultures of *P. aeruginosa* PAO1, PAO1ΔPf4, and PAO1ΔPilC were diluted to a concentration of 1×10^6^ CFU/mL in 5 mL LB broth. Aliquots of *P. aeruginosa* PAO1 were then infected Pf4 phage at MOIs ranging from 0.01-10. The samples were then incubated at 37°C shaking at 220 RPM for 18 hours to allow for superinfection. Then, all samples were set to an OD_600_ of 0.1 (2×10^7^ CFU/mL) in 1 mL of LB broth. Then, 150 µL was aliquoted to each well of a 96-well plate and incubated at 37°C non-shaking for 24 hours to allow for biofilms to form. Following the incubation, the well content was decanted, the wells were washed 2 times with DI water, and stained by adding 175 µL of 1% crystal violet solution followed by a 15 minute incubation at room temperature. The stain was removed from the wells, washed 3 times with DI water, and allowed to dry completely for several hours. After the plate was fully air dried, 200 µL of 30% acetic acid in water was added to each well to solubilize the crystal violet stain. The plate was incubated at room temperature for 15 minutes and the content of each well was transferred to a new 96-well plate, followed by measurement of absorbance at 550 nm using a Tecan M200Pro microplate reader. Wells containing only 30% acetic acid were used as a blank control.

### Macrophage Phagocytosis and Clearance Assays

#### General Procedure

The effect of SIE mechanisms on the phagocytosis and clearance of *P. aeruginosa* by macrophages was evaluated via the macrophage infection assay as previously described with some adaptations (52).

Overnight cultures of *P. aeruginosa* PAO1, PAO1ΔPf4, and PAO1ΔPilC were diluted to 1×10^6^ CFU/mL and a Pf4 phage was added to *P. aeruginosa* PAO1 at 1×10^7^ PFU/mL (MOI 10). The cultures were allowed to incubate for 18 hours, followed by a dilution to 2.5×10^6^ CFU/mL. A 200 µL aliquot of RAW 264.7 cells was added to each well of a 96-well plate at a density of 5×10^4^ cells/well in DMEM + 10% FBS and allowed to adhere overnight. Cell medium was removed and replenished with 200 µL of fresh DMEM containing 2.5×10^6^ CFU/mL of either *P. aeruginosa* PAO1, PAO1+Pf4 phage, PAO1ΔPf4, or PAO1ΔPilC for a MOI of 50. The plate was centrifuged at 500 × g for 30 minutes at 37°C to force macrophage infection. The plate was then incubated at 37°C + 5% CO_2_ for 30 minutes. After incubation, the cells were washed 3 times with phosphate buffered saline (PBS) and the media was replaced with DMEM + 100 µg/mL of gentamicin and incubated at 37°C + 5% CO_2_ for 1 hour to kill extracellular bacteria, resulting in a total infection time of 2 hours.

### Superinfection Phagocytosis Assay

Following incubation, cells were washed 3 times with PBS and lysed with 100 µL of 0.05% Triton X-100 (Promega). The contents of the well were serially diluted, plated on LB agar, and incubated at 37°C prior to CFU enumeration.

### Clearance Assay

Following the 2-hour incubation, cells were washed 3 times with PBS and some wells were lysed with 100 µL of 0.05% Triton X-100. Media in wells that were not lysed was replenished with DMEM + 25 µg/mL of gentamicin. These wells were allowed to incubate for an additional 2, 4, or 22 hours. They were then washed with PBS and lysed with 100 µg/mL of Triton X-100. At the end of each incubation and after cells were lysed, the contents of the well were serially diluted, plated on LB agar, and incubated at 37°C prior to CFU enumeration.

### Pf4 Phagocytosis Assay

An overnight culture of *P. aeruginosa* PAO1 was diluted to 2.5×10^6^ CFU/mL, then either left untreated or infected with Pf4 phage at a final concentration of 1×10^8^ PFU/mL. A 200 µL aliquot of RAW 264.7 cells was added to each well of a 96-well plate at a density of 5×10^4^ cells/well in DMEM + 10% FBS and allowed to adhere overnight. Cell medium was removed and replenished with 200 µL of untreated PAO1 or PAO1 infected with Pf4 phage at an MOI of 50. The plate was centrifuged at 500 × g for 30 minutes at 37°C to force macrophage infection. The plate was then incubated at 37°C + 5% CO_2_ for 30 minutes. After incubation, the cells were washed 3 times with phosphate buffered saline (PBS) and the media was replaced with DMEM + 100 µg/mL of gentamicin and incubated at 37°C + 5% CO_2_ for 1 hour to kill extracellular bacteria, resulting in a total infection time of 2 hours. Following incubation, cells were washed 3 times with PBS and lysed with 100 µL of 0.05% Triton X-100 (Promega). The contents of the well were serially diluted, plated on LB agar, and incubated at 37°C prior to CFU enumeration.

### Pseudomonas Phylogeny

Phylogenetic analysis of *P. aeruginosa* genomes was completed to identify the evolutionary relationships between strains containing or lacking Pf phages. A total of 405 *P. aeruginosa* whole genome sequences were obtained from the Pseudomonas Genome Database in FASTA format (26). The sequences were annotated using Prokka (53). The annotated genomes were aligned using Roary core genome alignment (54). The phylogenetic tree was built from this alignment using RAxML and was visualized using R studio, utilizing the ape package (55, 56).

### Pf Phage Identification

The presence of Pf phages in *P. aeruginosa* genomes was confirmed via nucleotide BLAST of each of the 405 genomes (57). The sequence of the *coaB* gene, the major coat protein of Pf phages, was obtained from the *P. aeruginosa* PAO1 reference strain in the Pseudomonas Genome Database and utilized to predict Pf phage presence as done previously (58). The BLASTN parameters were set to return sequences with ≥90% similarity and a query coverage of 100%.

### Phylogenetic Signal Analysis

The phylogenetic signal lambda was calculated as described previously utilizing the phylogenetic data and the presence or absence of the *coaB* gene as the designated trait (59). Two models were generated: One fitting the trait data to the phylogeny generated in this study, leaving lambda unconstrained, and another fitting the trait data to a “star” phylogeny with no structure, which constrains lambda to 0. The log-likelihoods of these models were then compared using a likelihood ratio test to determine the significance of the phylogenetic signal. Lambda values and p-values were generated for each case. All analyses were performed in R, including the caper and ape packages (56, 60).

### dN/dS Analysis

The analysis of codon sites of selection was performed using HyPhy and Fixed Effects Likelihood Modeling (FEL) and Mixed Effects Model of Evolution (MEME) (61–63). Briefly, alignments of *coaB* genes were generated using mafft, and were then passed through FEL and MEME analysis to identify sites of positive and purifying selection (64).

### Statistics

All statistical analyses were performed using GraphPad Prism 10.2.3. Statistical testing consisted of two-tailed Student’s t-tests, likelihood ratio tests, one-way ANOVAs, Welch’s ANOVAs, and two-way ANOVAs and significance levels were set at p<0.05. The necessary assumptions for each test, including normality, linearity, independence, and homogeneity of variance were checked. Any post-hoc analyses were performed utilizing Dunnett’s multiple comparisons test or Dunn’s multiple comparisons test where treatment groups were compared to an untreated control. All tests were run in triplicate unless otherwise stated. All graphs were generated using GraphPad Prism 10.2.3 unless otherwise stated.

## Acknowledgments

We would like to thank the Secor Lab at Montana State University for providing us with the *P. aeruginosa* PAO1ΔPf4 mutant strain. D.M.C. was in part supported by the University of Florida Opportunity Fund awarded to D.M.C. M.J.B. was partly supported by the University Scholars Honors Program, University of Florida.

